# An age-dependent reversal of the baseline-inflammation to vaccine-antibody-response relationship: a meta-analytic replication in ImmuneSignatures2 with independent influenza cohorts

**DOI:** 10.64898/2026.06.14.732119

**Authors:** Kilian Maire

## Abstract

**Background:** A pro-inflammatory transcriptional setpoint measured before vaccination predicts a stronger antibody response in younger adults but loses that value, or reverses, in older adults. This reversal has been characterised almost entirely for influenza, and it is unknown whether it is specific to influenza or a more general feature of immune ageing.

**Results:** Using the harmonised HIPC-II ImmuneSignatures2 compendium and two independent cohorts processed from raw data with the same pipeline, we built a baseline inflammation score from inflammatory and NF-kB blood transcription modules and tested its interaction with age on the antibody response. The inflammation-by-age interaction was negative in a pooled mixed model (t between −2.1 and −3.0 across three score definitions) and for a binarised high-responder outcome (p = 0.015). For influenza it survived leave-one-study-out, the non-normalised data, continuous-age and study-fixed-effect specifications. A random-effects meta-analysis of the old-minus-young inflammation-to-response slope difference across five cohorts, two of them independent, gave −0.15 (95% CI −0.27 to −0.03; p = 0.015; I-squared = 0%). An independent out-of-compendium replication across five Stanford influenza seasons (115 unique donors after deduplicating repeated enrolments) reproduced the negative interaction in direction in every season (pooled −0.23; I-squared = 0%), though its deduplicated interval still included zero. Within the compendium the reversal was present among viral vaccines (extending from influenza to a single small zoster cohort, 35 adults) and absent for recombinant hepatitis B; a formal vaccine-platform moderator was not significant (p = 0.47) and could not be separated from whether the response is primary or recall.

**Conclusions:** This confirmatory reanalysis replicates and formally quantifies a previously reported, age-dependent reversal of the baseline-inflammation benefit for the vaccine antibody response (Avey et al., Science Immunology 2017; Fourati et al., Nature Communications 2016). Its contribution is not the effect’s existence but its formal estimation as an inflammation-by-age interaction across the ImmuneSignatures2 compendium, together with an independent, out-of-compendium replication across five Stanford influenza seasons (consistent in direction in every season, I-squared = 0%, though the deduplicated interval still spanned zero). Whether the reversal generalises beyond influenza to other vaccine types could not be established: a systematic search of the public systems-vaccinology record (ImmuneSpace) found no adequately powered non-influenza cohort, so the cross-type question remains open, neither confirmed nor refuted.

## Background

Older adults respond less well to vaccination than younger adults, and the gap is a major public-health problem because the same people carry the highest burden of infectious disease. The decline is driven in large part by immunosenescence, the remodelling of the immune system with age that weakens the generation of protective antibody and raises susceptibility to infection [1, 2]. Improving vaccine protection in older adults requires understanding not only that responses fall, but why the rules that predict a good response in the young appear to change with age.

Beginning with early-signature analyses of yellow-fever and seasonal-influenza vaccination [3, 4], a large body of systems-vaccinology work has shown that the response is partly set before the vaccine is given: the baseline state of the immune system, read from blood gene expression, predicts how strongly a person will later respond [5, 6, 7]. One of the more reproducible baseline signals is a pro-inflammatory, NF-kB-driven setpoint (NF-kB is a central inflammatory signalling pathway). In younger adults a higher inflammatory tone before vaccination tends to accompany a stronger antibody response. Ageing, however, is itself accompanied by a chronic, low-grade inflammatory state, termed inflammaging, that is mechanistically entangled with immunosenescence [8, 9] and accumulates as a systemic, low-grade inflammatory burden across the lifespan [10, 11]. An immune system already running an inflammatory program may have little to gain, and may even be impaired, when asked to mount a fresh response.

Consistent with this, the predictive value of the inflammatory baseline is not fixed across the lifespan. In influenza cohorts a baseline inflammatory signature is positively associated with the antibody response in younger adults but the association inverts in older adults [6]. A subsequent pan-vaccine analysis of pre-vaccination endotypes restricted its primary analysis to adults aged 18 to 55 and noted that the inflammatory endotype was not predictive in older people for several vaccines [7], but it did not test an age-by-inflammation interaction formally, nor ask whether the inversion observed for influenza holds for other vaccines. Almost all of the evidence for the reversal comes from influenza, and older-adult systems-vaccinology data for other vaccines are scarce. The contribution here is therefore neither the existence of the influenza inversion, established by [6], nor the hepatitis B hyporesponse, reported by [17], but a formal age-by-inflammation interaction tested across vaccines with two independent replications processed from raw data, together with the recognition that the inflammatory predictor’s direction is confounded between vaccine platform and the primary-versus-recall nature of the response. This is therefore an explicitly confirmatory study: it does not supersede the original reports but strengthens them by quantifying the interaction formally and replicating it out-of-compendium, and by delimiting honestly where the present evidence stops.

We asked a single, falsifiable question: does the age-dependent reversal of the baseline-inflammation to antibody-response relationship extend beyond influenza? We used the harmonised ImmuneSignatures2 compendium [12, 13] and two independent cohorts processed from raw data with the same pipeline, fit the interaction directly, subjected it to a battery of robustness checks, and combined the evidence across vaccines (Fig. 1). A reversal confined to influenza would point to influenza-specific biology; one shared by other vaccines would implicate immune ageing more broadly; and a reversal seen for some vaccines but not others would suggest that the inflammatory predictor is conditional on vaccine type as well as age. Testing the interaction across several vaccines, rather than in influenza alone, is what lets these possibilities be told apart.

**Fig. 1.**
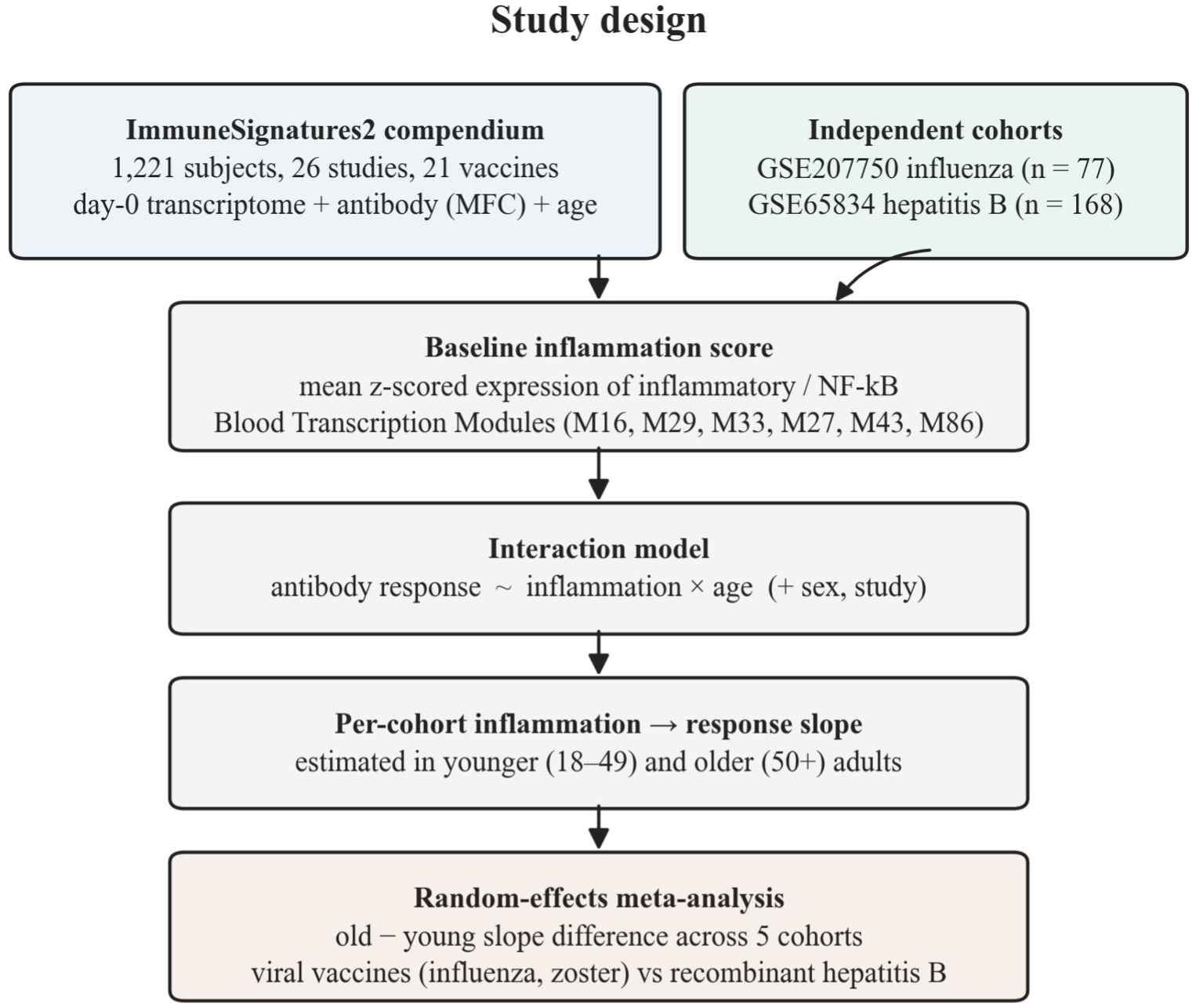
Study design. Day-0 transcriptomes, antibody responses and age from the ImmuneSignatures2 compendium and two independent cohorts feed a single baseline inflammation score; the inflammation-by-age interaction is estimated per cohort and combined by meta-analysis, with viral vaccines contrasted against recombinant hepatitis B.

## Methods

Data. The primary data were the HIPC-II ImmuneSignatures2 cross-study ExpressionSet [12, 13], which harmonises day-0 (pre-vaccination) blood transcriptomes with post-vaccination antibody responses, age, sex, vaccine and study for 1,221 subjects across 26 studies and 21 vaccines. Antibody response was the maximum fold change (MFC, the largest post-vaccination rise in antibody titre relative to the pre-vaccination day-0 level) provided by the resource. Subjects were grouped as younger (18 to 49 years) or older (50 years and above).

Baseline inflammation score. The inflammation score was the mean, over a pre-specified set of inflammatory and NF-kB Blood Transcription Modules [14] (predefined sets of co-expressed blood genes; here M16 TLR/inflammatory signalling, M29, M33 inflammatory response, M27, M43 and M86), of each gene’s z-scored day-0 expression. To guard against choosing a module set after seeing the result, three score definitions were registered in advance (the full inflammatory set, an NF-kB core and the single inflammatory-response module M33); all three are reported.

Statistical models. Because antibody assays differ across vaccines and cohorts (maximum fold change, log2 haemagglutination-inhibition titre fold change and log10 anti-HBs concentration), every response was standardised within its study or cohort before pooling, so the heterogeneous readouts enter the meta-analysis as slope differences on a common standardised scale rather than in their native units. The primary model was a linear mixed model, MFC ∼ inflammation_z * age_group + sex + (1 | study), with the inflammation-by-age interaction as the quantity of interest, fit pooled, influenza-only, influenza-removed and within each vaccine. Per-vaccine inflammation-to-response slopes were combined by random-effects meta-analysis, with heterogeneity summarised by the Q statistic, tau-squared and a 95% prediction interval and robustness checked by leave-one-cohort-out. Robustness of the pooled interaction was assessed with Satterthwaite p-values and confidence intervals (lme4 and lmerTest), a continuous-age parameterisation, the non-normalised expression matrix, leave-one-influenza-study-out refitting, a study fixed-effect specification, a comparison against random inflammation-slope-by-study models, a per-subject leave-one-out influence diagnostic, and a binarised high-responder outcome (the ImmuneSignatures2 high- versus low-responder classification, dropping the moderate class, fitted by a logistic mixed model with the same fixed and random terms). To check that the interaction was not a mechanical consequence of older adults responding within a narrower range, three further controls compared the standardised-response dispersion between age groups for each of the three age-complete vaccines, refitted the interaction after reweighting younger subjects to equalise the response variance of the two arms, and located the inflammatory-score interaction within an empirical null built from 1,000 random gene sets of the same size as the inflammatory set. A vaccine-platform moderator (viral versus recombinant protein), and the equivalent primary-versus-recall moderator, were tested by meta-regression, and the moderator’s statistical power was evaluated by simulation. As an exploratory check that the score was not driven by a single gene, the interaction model was also refit for each inflammatory-module gene separately and the consistency of the per-gene interaction signs summarised by a binomial sign test.

Independent cohorts. Two cohorts outside ImmuneSignatures2 were harmonised with the identical inflammation score and model. GSE207750 (University of Georgia seasonal influenza) [15] provided day-0 whole-blood RNA-seq; per-subject age and haemagglutination-inhibition titres (HAI, the standard influenza antibody assay) were taken from the open companion supplement [16] and joined by subject identifier (77 of 77 matched; ages 21 to 80; response was the maximum log2 titre fold change across the four vaccine strains). GSE65834 (Twinrix hepatitis B) [17] was self-contained, with pre-vaccination expression, age and post-vaccination anti-HBs titre per subject (anti-HBs is the antibody to the hepatitis B surface antigen, the protective response; 168 subjects, ages 25 to 83, 142 of them older adults; response was log10 anti-HBs concentration). Analyses used R with the lme4 [18] and metafor [19] packages.

Supplementary influenza replication (Stanford SLVP). As an additional, separately reported test of the influenza interaction, five trivalent influenza seasons of the Stanford Lewis-Sigler Vaccine Project (ImmuneSpace SDY112, SDY311, SDY312, SDY314, SDY315; none in ImmuneSignatures2) were assembled through the ImmuneSpace LabKey interface: day-0 expression matrices, HAI titres, age and the expression-sample-to-donor maps. Response was the maximum log2 HAI fold change (day 28 over day 0) across the season’s assayed strains, and the inflammation score used the same blood transcription modules, z-scored within each season’s baseline samples. Because the project is longitudinal, the same donors recur across seasons; they were identified by donor accession and deduplicated. The primary pooled fit retained every donor-season with study and donor as crossed random effects, with a one-season-per-donor fit as sensitivity. This series is reported outside the six-cohort meta-analysis because it falls outside the public, no-login eligibility gate and because its repeated-donor structure requires dedicated modelling.

Cohort eligibility. Candidate datasets were screened against six pre-specified mechanical gates applied identically to each: a per-subject day-0 bulk transcriptome with genome-wide coverage; that transcriptome being public with no login or data-access approval; a post-vaccination antibody titre that is public per subject and joinable to the transcriptome by subject identifier; per-subject age; a usable age signal, defined as a within-cohort older-versus-younger contrast for the binary meta-analysis or a real continuous age span for the continuous-age meta-analysis; and independence from ImmuneSignatures2 to avoid double-counting. A dataset entered only if it passed all applicable gates, and each excluded dataset is recorded against the single first gate it failed; the full pass and fail status of every screened cohort against each gate is given in the Additional file. Of 24 candidate datasets screened, 6 passed all applicable gates and were included (the influenza, zoster and hepatitis B strata of ImmuneSignatures2 and the three independent cohorts GSE207750, GSE65834 and GSE124533); the remaining 18 were excluded at their first failing gate (6 at the day-0 bulk-transcriptome gate, 1 at public access, 6 at a public per-subject antibody, 1 at per-subject age, 1 at a usable age signal and 3 for already being within ImmuneSignatures2; Fig. S1).

## Results

Subjects with both an age group and a response spanned influenza (504 younger, 361 older), hepatitis B (25 younger, 135 older) and zoster (16 younger, 19 older); other non-influenza vaccines in the compendium had few or no older adults (Fig. 2A). Throughout, a recall response boosts pre-existing immunity (seasonal influenza; Zostavax boosting latent varicella-zoster memory) and a primary response is mounted against a neoantigen with no relevant pre-existing immunity (hepatitis B in naive adults; yellow fever in flavivirus-naive adults). Table 1 sets out the vaccines by platform and by response type.

**Fig. 2.**
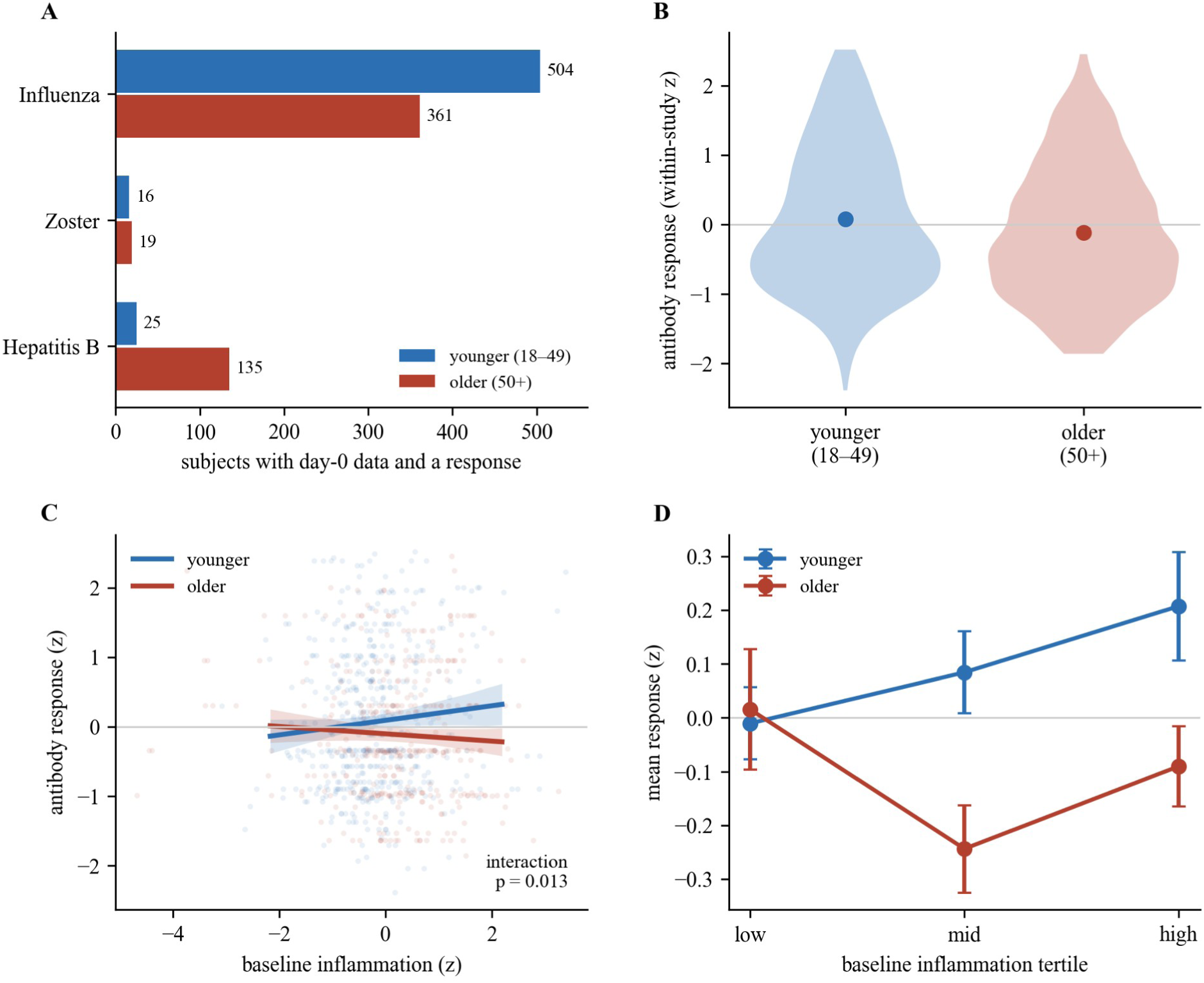
Baseline inflammation, age and the influenza antibody response. (A) Subjects with day-0 data and a measured response, by vaccine and age group. (B) Standardised antibody response by age group. (C) Response against baseline inflammation, with fitted lines and bootstrap confidence bands for younger (blue) and older (red) adults; the lines cross. (D) Mean response across inflammation tertiles by age group.

**Table 1.** Vaccines analysed, by platform and by whether the antibody response is a recall of pre-existing immunity or a primary response to a neoantigen, and whether the cohort has an older-adult arm. The viral/recombinant and recall/primary axes coincide for every vaccine except yellow fever (viral but a primary response), the only cohort that could separate them, which has no older-adult arm; this is why the two framings cannot be distinguished here (Fig. 5B).

| Vaccine | Platform | Response type | Older-adult arm | In analysis |
| --- | --- | --- | --- | --- |
| Influenza (TIV) | viral, inactivated | recall | yes | binary + continuous |
| Zoster (Zostavax) | viral, live attenuated | recall | yes | binary + continuous |
| Hepatitis B | recombinant protein | primary | yes | binary + continuous |
| Yellow fever | viral, live attenuated | primary | no (younger only) | continuous only |
| Meningococcus | conjugate / polysaccharide | primary | no (younger only) | continuous only |

Baseline inflammation interacted with age on the antibody response. Older adults responded less well overall (Fig. 2B), and in the pooled mixed model the inflammation-by-age interaction was negative across all three inflammation-score definitions (t between −2.1 and −3.0); a binarised high-responder outcome agreed (pooled logistic p = 0.015). Within influenza, inflammation predicted a higher response in younger adults (slope +0.11) and a lower response in older adults (slope −0.06), with an interaction of −0.17 (t = −2.5, p = 0.013; Fig. 2C). The same crossover was visible as a monotone rise of response across inflammation tertiles in the young that flattened and reversed in the old (Fig. 2D).

Within ImmuneSignatures2, a meta-analysis of the old-minus-young slope difference across influenza, zoster and hepatitis B gave −0.16 (95% CI −0.29 to −0.04; p = 0.012; I-squared = 0%), with zoster moving in the same direction as influenza and hepatitis B flat (Fig. 3A). The two independent cohorts reproduced the pattern by vaccine type (Fig. 3B): the independent influenza cohort showed the same crossover (younger slope +0.03, older slope −0.03, difference −0.06), while the independent hepatitis B cohort, with 142 older adults, showed no reversal (continuous interaction −0.005, p = 0.92; slope difference −0.04). Hepatitis B therefore showed no reversal in two independent cohorts. Both, however, are weighted toward older adults (25 versus 135, and 26 versus 142), so the younger-adult slope that creates the crossover is estimated from relatively few subjects in each; the consistent hepatitis B null is best read as a reproducible absence of signal rather than as a precisely estimated zero.

**Fig. 3.**
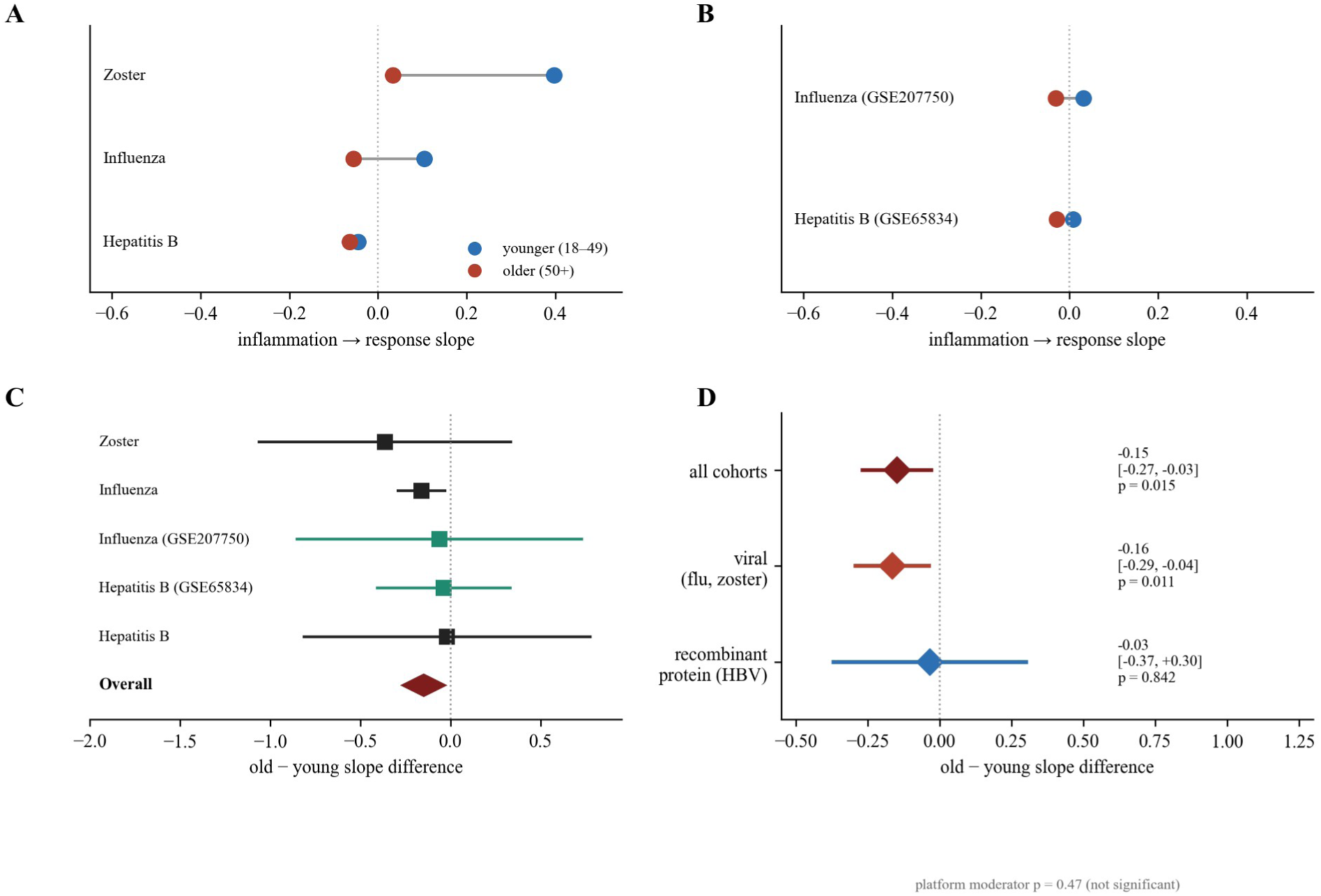
Generalisation across vaccines and cohorts. (A) Inflammation-to-response slope in younger (blue) and older (red) adults for each ImmuneSignatures2 vaccine. (B) The same for the two independent cohorts. (C) Random-effects meta-analysis of the old-minus-young slope difference across five cohorts (independent replications in green; diamond, overall estimate). (D) Subgroup estimates for all cohorts, viral vaccines and recombinant hepatitis B.

The independent influenza signal was probed further in the Stanford Lewis-Sigler Vaccine Project, five trivalent inactivated influenza seasons (ImmuneSpace SDY112, SDY311, SDY312, SDY314 and SDY315), each an older-versus-younger Fluzone cohort outside ImmuneSignatures2 and analysed with the identical inflammation score and model. These are reported separately from the primary meta-analysis because the resource requires registration (outside the public, no-login eligibility gate) and, being longitudinal, re-enrols the same donors across seasons: the 371 donor-season records map to only 115 unique donors (45 younger, 70 older), so repeated enrolments were deduplicated on donor identity and modelled as a random effect. The inflammation-by-age interaction was negative in direction in all five seasons with no detectable heterogeneity (I-squared = 0%); the pooled estimate from a mixed model over all donor-seasons (younger and older slope contrast, donor and season as random effects) was −0.23 (95% CI −0.57 to +0.11), and a sensitivity fit keeping one season per donor gave −0.32 (95% CI −0.92 to +0.28). The point estimate matches the ImmuneSignatures2 influenza interaction (−0.17), but once the repeated-donor structure is accounted for the confidence interval still spans zero: the series is roughly 45 unique younger donors, so it corroborates the direction of the reversal across five independent seasons without, on its own, providing a decisive independent confirmation (Fig. S2).

Combining all five cohorts, two of them independent, the random-effects meta-analysis of the old-minus-young slope difference was −0.15 (95% CI −0.27 to −0.03; p = 0.015; Fig. 3C), a small effect. Heterogeneity was not detectable (Q = 0.9 on 4 df, p = 0.93; tau-squared = 0; I-squared = 0%) and the 95% prediction interval (−0.27 to −0.03) also excluded zero, though with five cohorts heterogeneity is estimated too imprecisely to be read as positive evidence of consistency. The estimate survived removal of any single cohort except the main influenza stratum (leave-one-cohort-out; dropping influenza left −0.09, p = 0.52), so it is carried by influenza; removing zoster left −0.14 (p = 0.02), but because zoster is the only non-influenza viral cohort the generalisation beyond influenza itself rests on that single cohort.

The viral-vaccine subgroup (influenza twice and zoster) was −0.17 (p = 0.011) and the recombinant-protein subgroup (hepatitis B twice) was −0.03 (p = 0.84); a formal vaccine-platform moderator was not significant (p = 0.47). That non-significance is uninformative rather than evidence of similarity: with two platform groups and imprecise hepatitis B slopes the moderator has very low power (about 10% for the observed difference; Discussion), so it neither establishes nor excludes a platform difference (Fig. 3D).

The influenza interaction was robust. It stayed negative on both the re-normalised and the raw, non-normalised expression matrix (Fig. 4A), held across the three inflammation-score definitions (Fig. 4B), persisted under a continuous-age model (Fig. 4C), and survived removal of any single study (sixteen leave-one-study-out estimates, all negative, range −0.23 to −0.13; Fig. 4D); a study fixed-effect specification gave an identical estimate (−0.167, p = 0.017). Allowing the inflammation slope to vary by study (a random-slope model) left the interaction negative and if anything larger and was not favoured over the random-intercept model (likelihood-ratio p = 0.14), so the random-intercept specification is not anti-conservative here; and no single subject drove the result (largest leave-one-subject-out change 11% of the estimate with the sign unchanged, and dropping the most influential 1% of subjects left −0.15). Pooling each cohort’s continuous-age interaction by vaccine platform, the viral subgroup remained negative (three age-informative cohorts, −0.08, p = 0.03) whereas the recombinant-protein subgroup did not (two cohorts, −0.005, p = 0.91); adding the age-incomplete cohorts that carry near-zero weight left this unchanged (Fig. 5A). Regrouping the same cohorts by whether the vaccine elicits a recall or a primary response, rather than by platform, gave a numerically identical contrast (recall −0.08, p = 0.03; primary −0.005, p = 0.91; Fig. 5B), because the only cohort that changes group, yellow fever, has no older-adult arm and so carries near-zero weight.

**Fig. 4.**
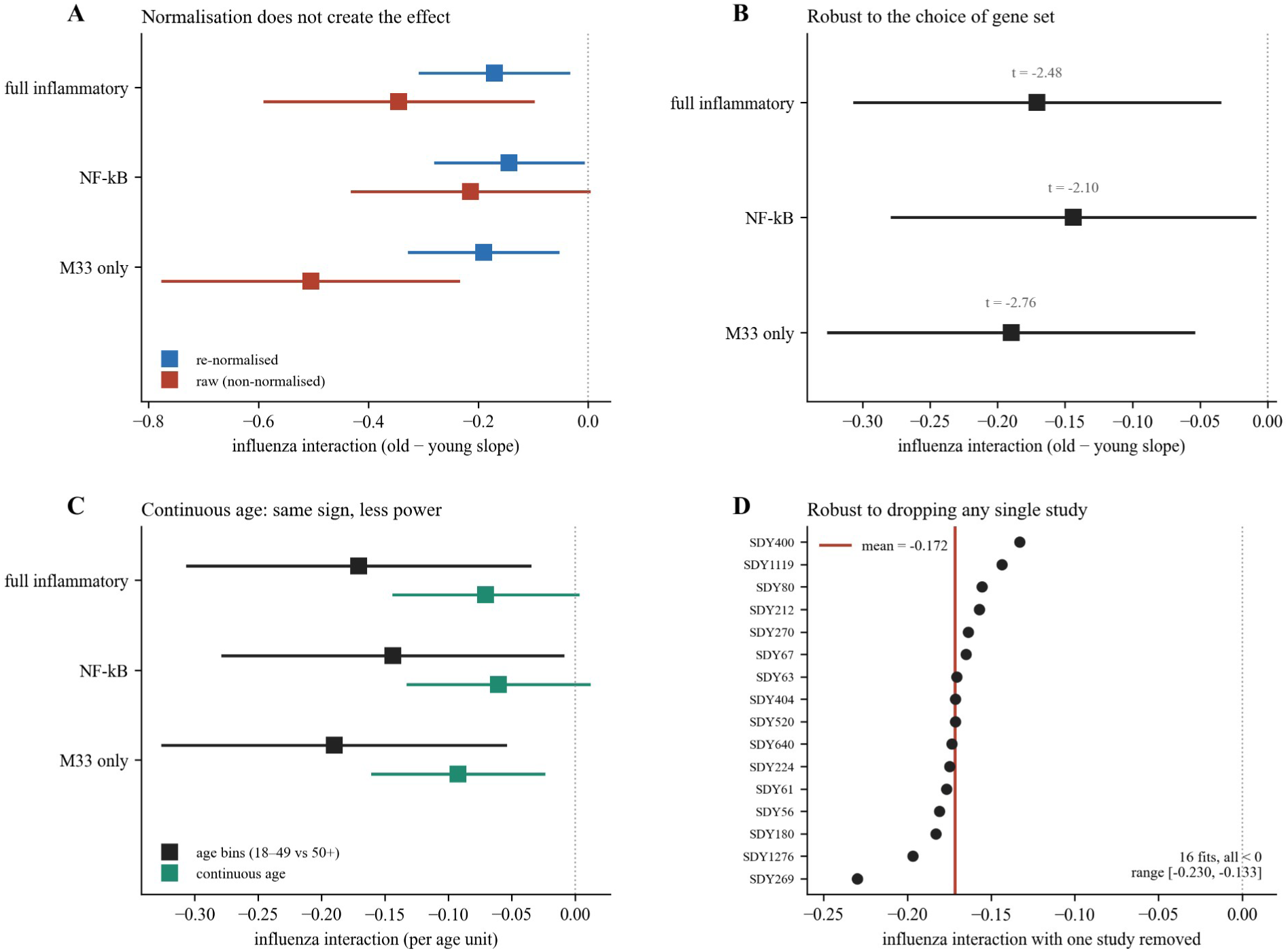
Robustness of the influenza inflammation-by-age interaction. (A) Re-normalised versus raw (non-normalised) expression, for each inflammation score. (B) Sensitivity to the inflammation gene set (full, NF-kB core, M33 only), with t values. (C) Dichotomised age (18 to 49 versus 50 and over) versus continuous age. (D) Leave-one-study-out re-fits across the sixteen influenza studies, all negative.

**Fig. 5.**
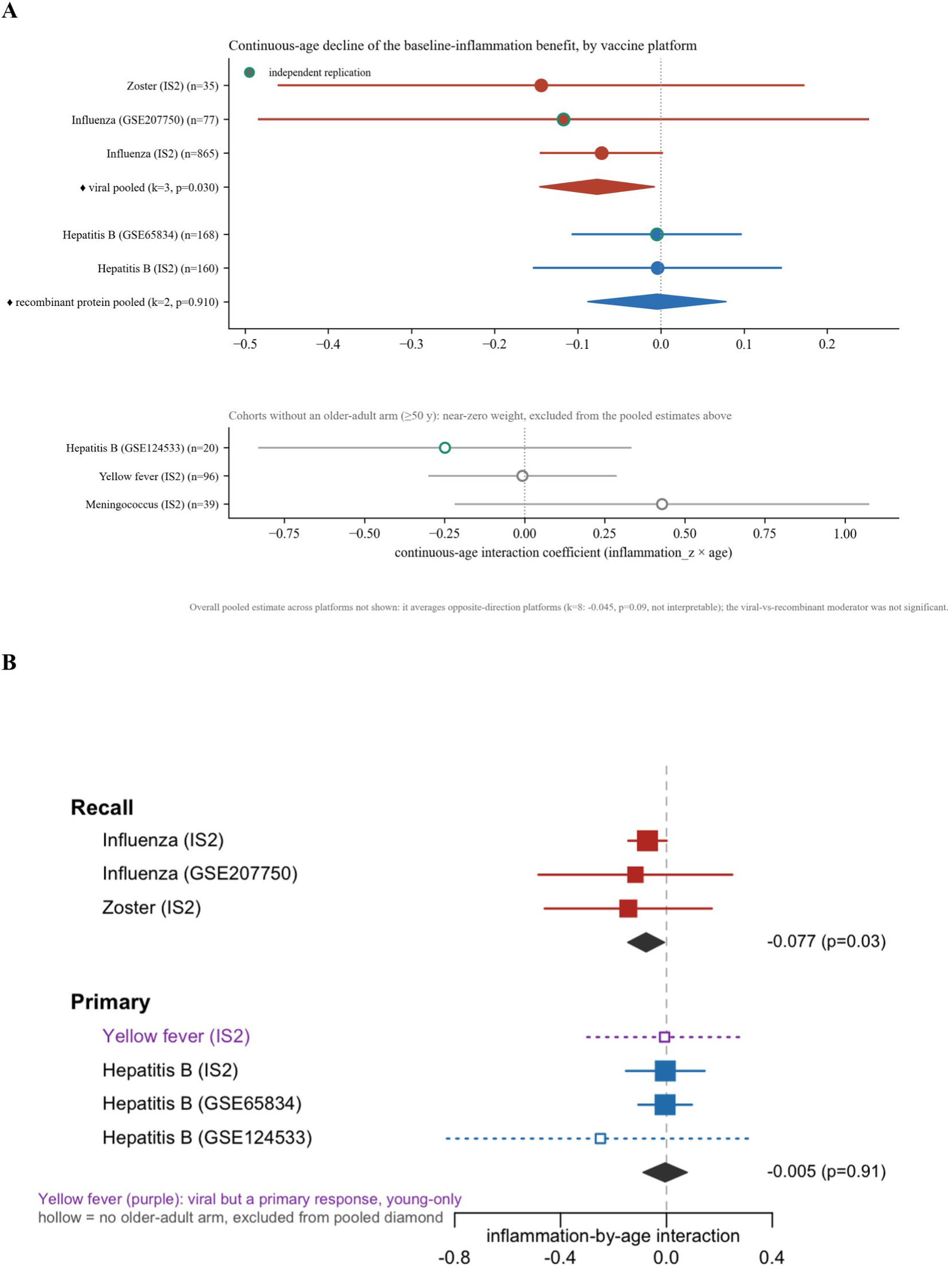
Continuous-age per-cohort inflammation-by-age coefficient. (A) Grouped by vaccine platform: cohorts with an older-adult arm and the per-platform pooled estimates (diamonds, independent replications outlined in green); cohorts without an older-adult arm, which carry near-zero weight and cannot inform an age reversal, are shown in a separate strip below. The viral pooled estimate is negative and the recombinant-protein estimate is null. (B) The same cohorts regrouped by whether the vaccine elicits a recall or a primary response. Filled markers have an older-adult arm and contribute to the pooled diamond; hollow markers are young-only and are excluded. The one cohort that changes group, yellow fever (purple, a viral vaccine giving a primary response), is young-only, so the recall and primary diamonds are identical to the viral and recombinant-protein ones in (A): the platform and primary-versus-recall framings cannot be separated on these data.

The interaction was not an artefact of a narrower response range in older adults. Within influenza the older arm’s standardised-response SD was only 7% smaller than the younger arm’s (ratio 0.93; variance-equality F-test p = 0.14), and reweighting subjects to equalise the two arms’ variances left the interaction essentially unchanged (−0.170, t = −2.5, versus −0.171). Arm-specific slopes crossed sign (younger +0.11, older −0.06) rather than shrinking in proportion to variance, which range compression alone cannot produce. The same control is less clean for the non-influenza cohorts: in both hepatitis B and zoster the older arm’s response variance is far smaller than the younger arm’s (variance ratios 0.14 and 0.10; F-test p < 0.001), driven largely by their small, imbalanced younger arms, and equalising the two arms’ variances moves the hepatitis B interaction slightly more negative (−0.02 to −0.07) and roughly halves the zoster interaction (−0.29 to −0.14). These variance-equalised refits are themselves unstable because they divide by a younger-arm SD estimated from about 25 subjects, so they do not establish a masked hepatitis B effect; rather, they show that the non-influenza estimates are sensitive to the variance treatment and too imprecise to support a strong platform contrast, consistent with the low power of the formal moderator. Against an empirical null of 1,000 same-size random gene sets the inflammatory score’s interaction fell at the 7.8th percentile (two-sided p = 0.08); it was therefore more extreme than 92% of random sets, though the null was itself mildly shifted toward negative interactions (mean −0.06), so this placebo test is directionally supportive without reaching conventional significance.

Decomposing the influenza score gene by gene confirmed that the interaction was broadly signed across the inflammatory set, most strongly in the inflammatory-response (M33) and TLR-signalling (M16) modules, rather than a single-gene effect. Of 106 inflammatory-module genes fitted individually, 74 (70%) carried a negative interaction (sign test p = 3 × 10⁻⁵), although few reached significance alone, as expected when each gene carries the shared effect attenuated by its own measurement noise. The same decomposition showed no directional consistency for hepatitis B, in ImmuneSignatures2 (56 of 106 negative; sign test p = 0.31) and again in the independent, older-weighted GSE65834 cohort (58 of 122 negative; sign test p = 0.74), so the coherence is specific to the influenza data and not an artefact of the score (Fig. 6).

**Fig. 6.**
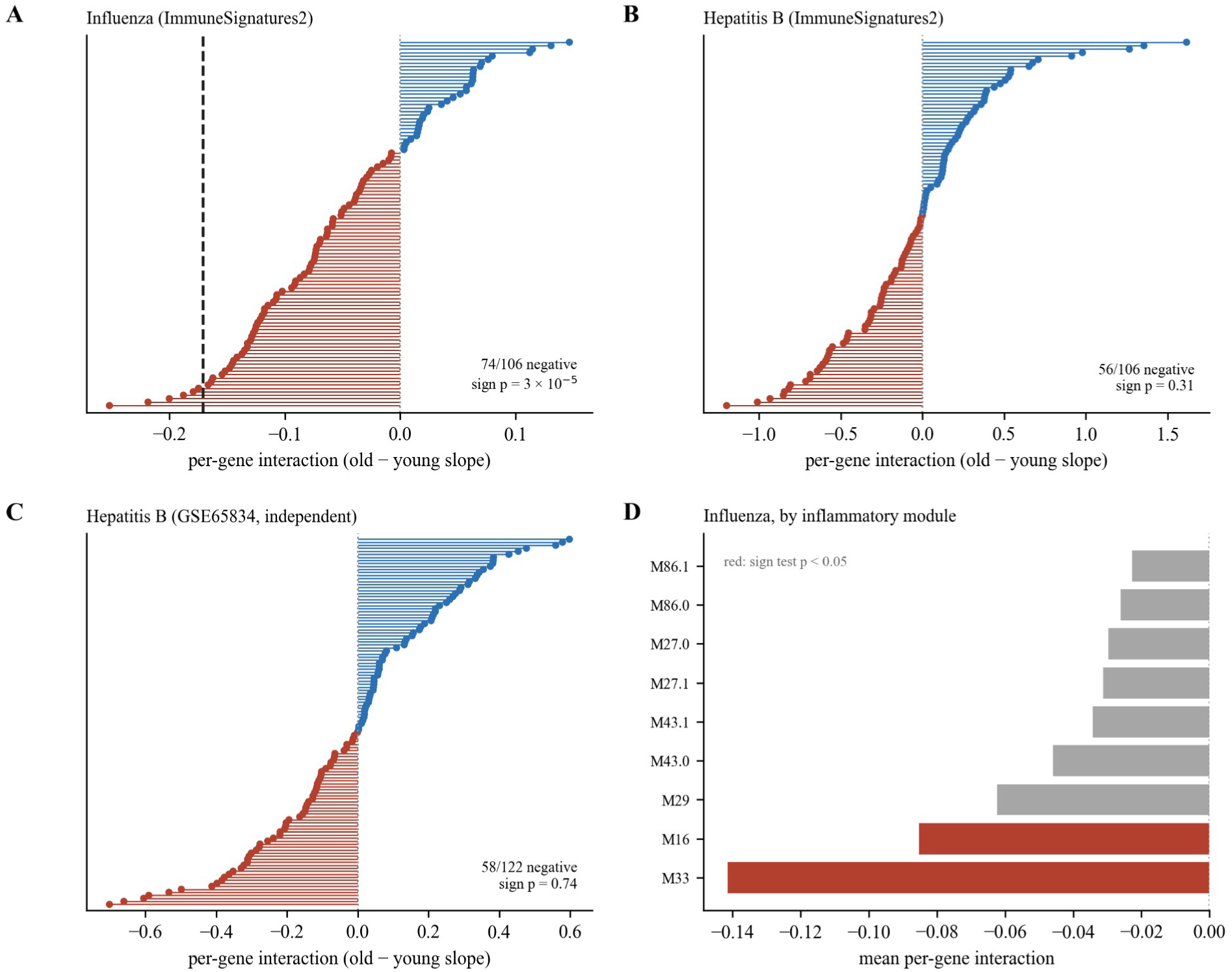
Gene-level decomposition of the interaction. (A) Influenza, each inflammatory-module gene fitted individually; most genes are negative (sign test p = 3 × 10⁻⁵), the dashed line marks the module-score estimate. (B) Hepatitis B in ImmuneSignatures2 and (C) the independent GSE65834 cohort show no directional consistency. (D) Influenza per-gene interaction averaged within each inflammatory module; red marks modules with a sign test p < 0.05.

## Discussion

The reversal of the baseline-inflammation to antibody-response relationship with age is not specific to influenza. It reproduces for zoster (Zostavax, a live attenuated vaccine; ImmuneSignatures2 study SDY984), a second viral vaccine, and the combined estimate across five cohorts, including two independent replications, is significant and homogeneous. Equally clearly, the reversal does not reach every vaccine. The recombinant hepatitis B vaccine showed no age reversal in two independent cohorts, one of them weighted toward older adults, and that null is a finding rather than a gap.

The hepatitis B result is consistent with that cohort’s own original report, which framed pre-vaccination inflammation as a marker of age-related hyporesponse rather than a benefit that reverses [17]. Read together, the cohorts suggest that an inflammatory baseline helps younger adults respond to live and inactivated viral vaccines and stops helping, or hinders, older adults, while for a recombinant protein antigen the inflammatory baseline carries little directional information at any age. A platform-dependent mechanism is an attractive summary, but we do not overstate it: the formal platform moderator was not significant, so the difference between the viral and hepatitis B vaccines is supported by two concordant hepatitis B cohorts rather than established statistically. The moderator is in fact severely underpowered: for the observed between-platform difference a five-cohort design of this composition has only about 10% power, and simulations indicate that no realistically attainable design, up to eight cohorts per platform with 150 subjects per age arm, would reach 80% (simulation, Methods). The platform question therefore cannot be settled by meta-analysis of per-cohort slope differences and would need a coordinated prospective study.

A platform account is, moreover, not the only reading of these data and may not be the most parsimonious. Among the three vaccines with both age groups, the viral-versus-recombinant contrast is the same partition of the data as a primary-versus-recall one: influenza and zoster both boost pre-existing immunity, whereas hepatitis B is a primary response to a neoantigen. The two moderators are therefore statistically indistinguishable here and return an identical estimate (p = 0.48). They diverge for only one cohort, yellow fever, a viral vaccine that elicits a primary response in naive adults: a platform account predicts a reversal for it, yet its inflammation-by-age interaction is flat (coefficient −0.01), which is instead what the primary-versus-recall account predicts. Consistent with this, when the continuous-age cohorts are grouped by response type the recall subgroup carries the reversal (−0.08, p = 0.03) and the primary subgroup does not (−0.005, p = 0.91), a contrast indistinguishable from the platform one (Fig. 5B). Yellow fever contributes only a within-younger age span, so this adjudicates weakly rather than decisively, but it raises the possibility that the operative variable is the nature of the response, recall versus primary, rather than the vaccine platform as such. Neither framing is established, and the two are confounded across the available vaccines; separating them requires cohorts that break their alignment, such as a primary-response viral vaccine or a recall recombinant-protein booster studied across the age range.

The practical implication is modest but useful. Baseline inflammatory signatures are sometimes proposed as general predictors of vaccine response, yet a transcriptional atlas of thirteen vaccines found no single universal signature of the antibody response [20]. These results add an age dimension to that caution, consistent with recent syntheses of immune ageing and vaccine response in older adults [21]: the predictive direction is conditional on both age and vaccine, and transporting a predictor trained in younger influenza vaccinees to older adults, or to a recombinant protein vaccine, is not safe. Consistent with a context-dependent role for the inflammatory baseline, inflammaging markers did not predict the waning of SARS-CoV-2 mRNA booster responses in a separate cohort, which tracked instead with markers of T-cell dysfunction [22].

This study has limits. The vaccine platforms with both age groups were influenza, zoster and hepatitis B; other older-adult non-influenza systems-vaccinology cohorts with a true day-0 baseline transcriptome, a per-subject antibody response and age are scarce. A systematic programmatic search of the ImmuneSpace catalogue (all studies, then those with a transcriptome, then their antibody assays) confirmed this directly: every non-influenza vaccine cohort with the required design is already inside ImmuneSignatures2, and the only independent non-influenza vaccine cohort outside it with a baseline transcriptome and an antibody readout (a zoster cohort) carried about ten subjects with a paired titre, far too few to test the interaction. The cross-type question is thus limited by a ceiling on the available public evidence, not resolved against the effect. Cohort eligibility was fixed by pre-specified mechanical gates applied identically to every candidate (Methods; Additional file, cohort screening table): the candidate yellow-fever dataset was excluded because no per-subject antibody titre is public and joinable to the transcriptome, and the pneumococcal dataset because its transcriptome is available only under controlled access, rather than by any judgement about the cohorts. Zoster rests on a single small cohort of 35 adults, which is the narrowest support in the study. The two independent cohorts each carry caveats: the influenza cohort’s antibody and age were joined from a companion supplement, and the hepatitis B cohort uses a different microarray platform. Baseline inflammation is a transcriptional module score, not a measured cytokine, and the platform moderator is underpowered (about 10% power). For influenza the interaction is robust to range compression and to variance matching and to subject-level influence, but in the much smaller, age-imbalanced hepatitis B and zoster cohorts the two age arms have very unequal response variance, so their estimates are sensitive to the variance treatment and should be read cautiously. The influenza interaction was further corroborated, in direction, across five independent Stanford SLVP seasons with no heterogeneity and a pooled estimate matching the main influenza value; that series is underpowered on its own, however, because its longitudinal design reduces roughly 400 records to about 115 unique donors, so its deduplicated interval still includes zero and it strengthens rather than settles the independent influenza case. Antibody readouts also differ in kind across vaccines (a fold change for influenza and zoster, an endpoint concentration for hepatitis B); within-study standardisation places them on a common scale but does not make them the same quantity, so the hepatitis B null could in part reflect its readout. Finally, the score’s specificity to the inflammatory gene set, tested against a random-gene-set null, was supportive in direction without reaching conventional significance, so a contribution from a non-specific transcriptional age signature cannot be excluded.

## Conclusions

The age-dependent loss of the baseline-inflammation benefit to vaccination, first reported for influenza, is here replicated and quantified formally as an inflammation-by-age interaction across the ImmuneSignatures2 compendium and corroborated, in direction, by an independent out-of-compendium replication across five Stanford influenza seasons. Within the compendium the reversal is also present in the viral stratum and absent for recombinant hepatitis B, but the non-influenza evidence is too thin, and the public record too sparse, to establish whether the reversal generalises across vaccine types; that question remains open. The practical message is the durable one: baseline inflammatory predictors of vaccine response should be treated as conditional on age rather than as universal signatures, and transporting a predictor trained in younger influenza vaccinees to older adults is not safe.

## Declarations

## Ethics approval and consent to participate

Not applicable. The study is a secondary analysis of publicly available, de-identified data; no new human or animal data were collected.

## Consent for publication

Not applicable.

## Availability of data and materials

All data are public. The ImmuneSignatures2 resource is at figshare (https://doi.org/10.6084/m9.figshare.17096978). The independent cohorts are in the Gene Expression Omnibus under accessions GSE207750, GSE65834 and GSE124533; per-subject antibody and age tables were drawn from the cited supplements. All analysis and harmonisation code, with a one-command reproduction (run_all.sh), is in a public Git repository (https://github.com/KilianMaire/vaccine-inflammation-age), with a versioned release archived on Zenodo (DOI 10.5281/zenodo.20688286).

## Additional files

Additional file 1: cohort screening table (mechanical inclusion and exclusion gates for every screened cohort). Additional file 2: random-gene-set null distribution for the inflammation-by-age interaction.

## Authors’ contributions

K.M. designed the study, performed the analyses and wrote the manuscript.

## Acknowledgements

This work used public data generated by the Human Immunology Project Consortium and by the investigators of the cited cohorts, whom we thank.

## Supplementary Figures

**Fig. S1.**
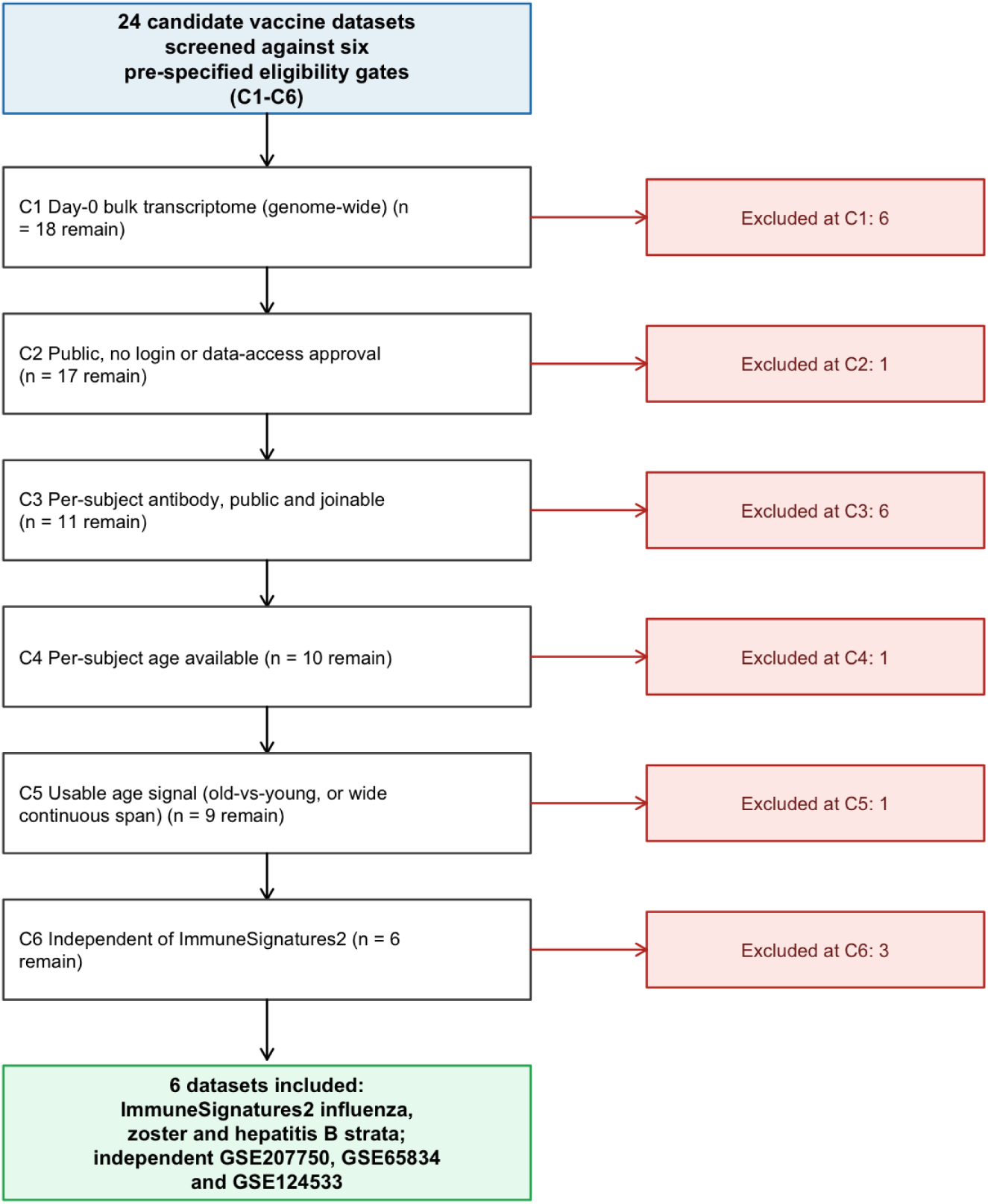
PRISMA-style flow of the mechanical cohort screen. Twenty-four candidate vaccine datasets were screened against six pre-specified eligibility gates applied in order (C1-C6); each excluded dataset is counted against the single first gate it failed. Six datasets passed all applicable gates and were included.

**Fig. S2.**
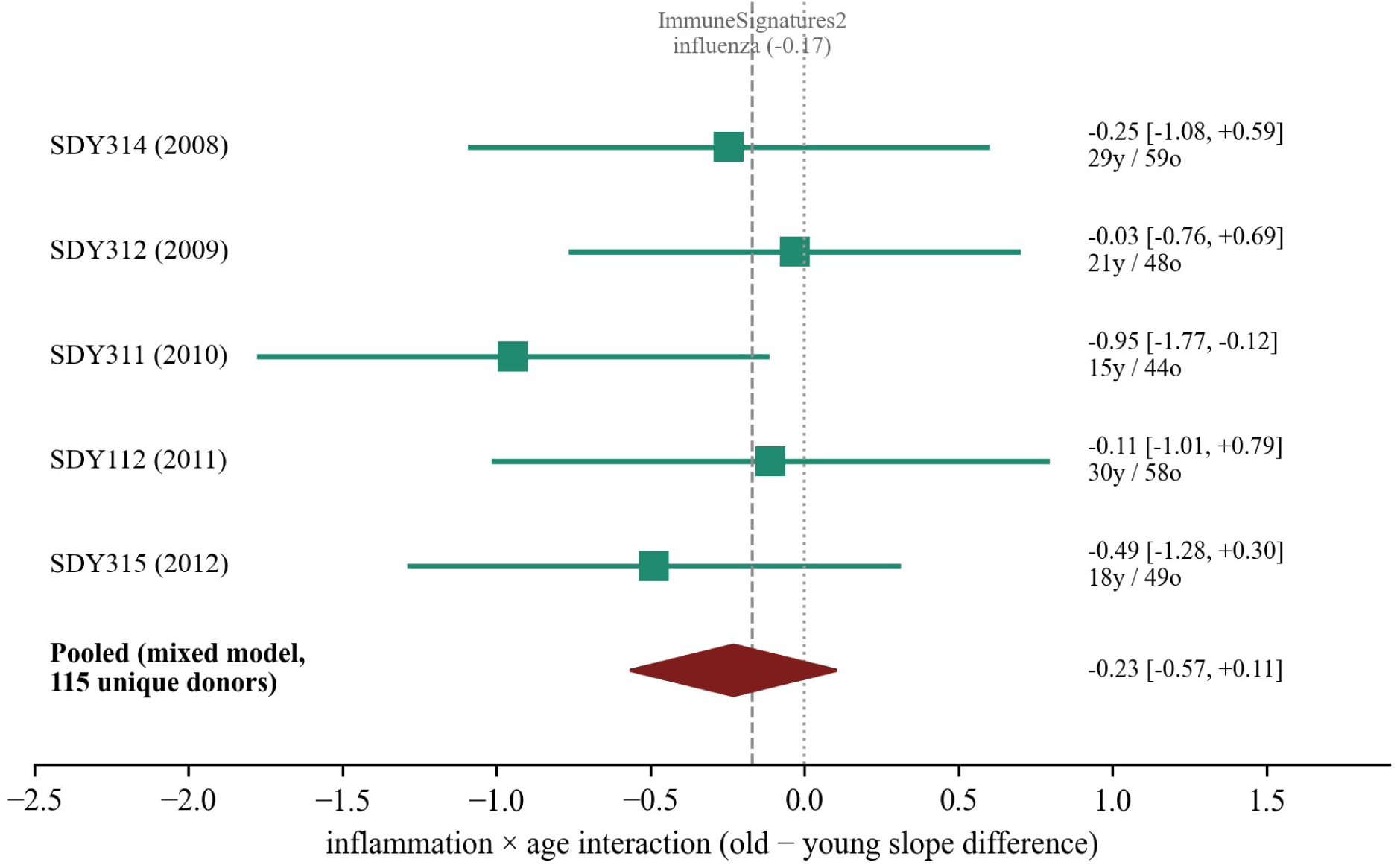
Supplementary independent influenza replication across the Stanford Lewis-Sigler Vaccine Project. Old-minus-young inflammation-to-response slope difference for each of five trivalent influenza seasons (ImmuneSpace SDY112, SDY311, SDY312, SDY314, SDY315; green squares, 95% confidence intervals; younger and older counts at right), with the pooled estimate from a mixed model over all donor-seasons after deduplicating to 115 unique donors (diamond). The dashed line marks the ImmuneSignatures2 influenza interaction (−0.17). The negative interaction is reproduced in direction in every season with no heterogeneity, but the deduplicated pooled interval still includes zero.

